# *Arabidopsis* MLO4 Functions as a Ca^2+^ Channel Essential for Mechanosensing in Root Tips

**DOI:** 10.1101/2022.06.05.494847

**Authors:** Zhengli Zhang, Yujia Sun, Pengfei Lu, Changxin Feng, Qi Niu, Geng Lin, Dongdong Kong, Liangyu Liu, Sheng Luan, Legong Li, Congcong Hou

## Abstract

Mechanical stimuli guide root growth to avoid obstacles in the soil. Although mechanisms underlying root mechanosensing remain largely unknown, calcium signaling is proposed to be an early event that proceeds directional root growth. In response to mechanical perturbations, root cells produce transient spikes in the cytosolic free Ca2+ levels. Loss-of-function mutations of MLOs cause root curling upon contact with hard surface of obstacles, the mechanism for MLOs action in root thigmomorphogenesis has been enigmatic. WT and mlo4 seedlings were grown on MS media containing gradient agar concentrations mimicking different levels of soil hardness and compared their root growth into the agar gradient mlo4 had much less roots penetrated the harder agar barrier as compared to WT roots. The channel activity of MLO4 was examined in Xenopus oocytes using the two-electrode voltage clamp (TEVC) method. The TEVC recording showed that the oocytes injected with MLO cRNA displayed large inward currents in the presence of Ca2+. MLO4 rescued the cch1/mid1 from growth arrest by the pheromone, these results indicate that MLO4 constitutes a novel Ca2+-permeable channel. Ca2+ spike was hardly detectable in mlo4/GCaMP6s seedings in response to a hard surface during root elongation. In summary, MLO4 functions as a typical Ca2+ channel that links touch stimulation to Ca2+ elevation in root tip cells.

## INTRODUCTION

Mechanical stimuli guide root growth to avoid obstacles in the soil. Although mechanisms underlying root mechanosensing remain largely unknown, calcium signaling is proposed to be an early event that proceeds directional root growth. In response to mechanical perturbations, root cells produce transient spikes in the cytosolic free Ca^2+^ levels [1,2]. Such Ca^2+^ elevations may be mediated by Ca^2+^-permeable channels directly or indirectly activated by mechano-stimuli. Several studies identified MSLs/CSCs/PIEZOs as mechanosensitive channels, but little is known about their function in mechanosensing of root tips. An intriguing study shows that loss-of-function mutations in several *MLOs* (*MILDEW RESISTANCE LOCUS Os*), including *MLO4* and *MLO11*, cause root curling upon contact with hard surface of obstacles, which are affected by auxin, pH and exogenous calcium levels [5]. Because biochemical function of MLO family proteins remained unknown for decades, the mechanism for MLO action in root thigmomorphogenesis has been enigmatic. Since all MLOs feature 6-7 predicted transmembrane domains [4], we hypothesized that MLO4 and MLO11 may act as calcium channels involved in root tip mechanosensing.

## RESULTS

We isolated a T-DNA insertion line (SAIL_266_C02) that lacked full-length transcripts of *MLO4* gene as measured by RT-PCR, suggesting that *mlo4* was a knock-out mutant allele (Supplementary Figure 1A-B). Based on previous study [5], *mlo4* displayed a touch-induced root curling phenotype. We decided to confirm this phenotype before using the mutant for other experiments. We grew WT and *mlo4* seedlings on MS media containing gradient agar concentrations mimicking different levels of soil hardness and compared their root growth into the agar gradient. For WT and *mlo4* mutant grown vertically at a 90 degrees right angle on the MS media containing 0.6% and 1.2% (2 mm thicker than upper) agar layers, first their roots all penetrated the 0.6% agar and then reached the surface of harder medium with 1.2% agar within 5 day after germination. After that only 22% of *mlo4* roots penetrated the harder agar barrier as compared to 93% of WT roots. In addition, *mlo4* exhibited root curling at the surface of hard barrier (Figure 1A-B). The barrier agar concentration was increased to 1.5%, the penetration percentage for WT was 87% while 23% of *mlo4* roots penetrated, however, both of WT and *mlo4* root had only 6% roots penetrated at a 1.75% barrier (Supplementary Figure 1C-F). A previous study shows that exogenous application of Ca^2+^ suppresses the root curling phenotype of *mlo4* [5]. Indeed, when we included 10 mM CaCl_2_ in the 1.2% agar medium, 85% of *mlo4* roots penetrated the barrier, similar to WT roots (Figure 1B), suggesting that MLO4 may be functionally important in Ca^2+^-mediated processes.

**Figure 1.**
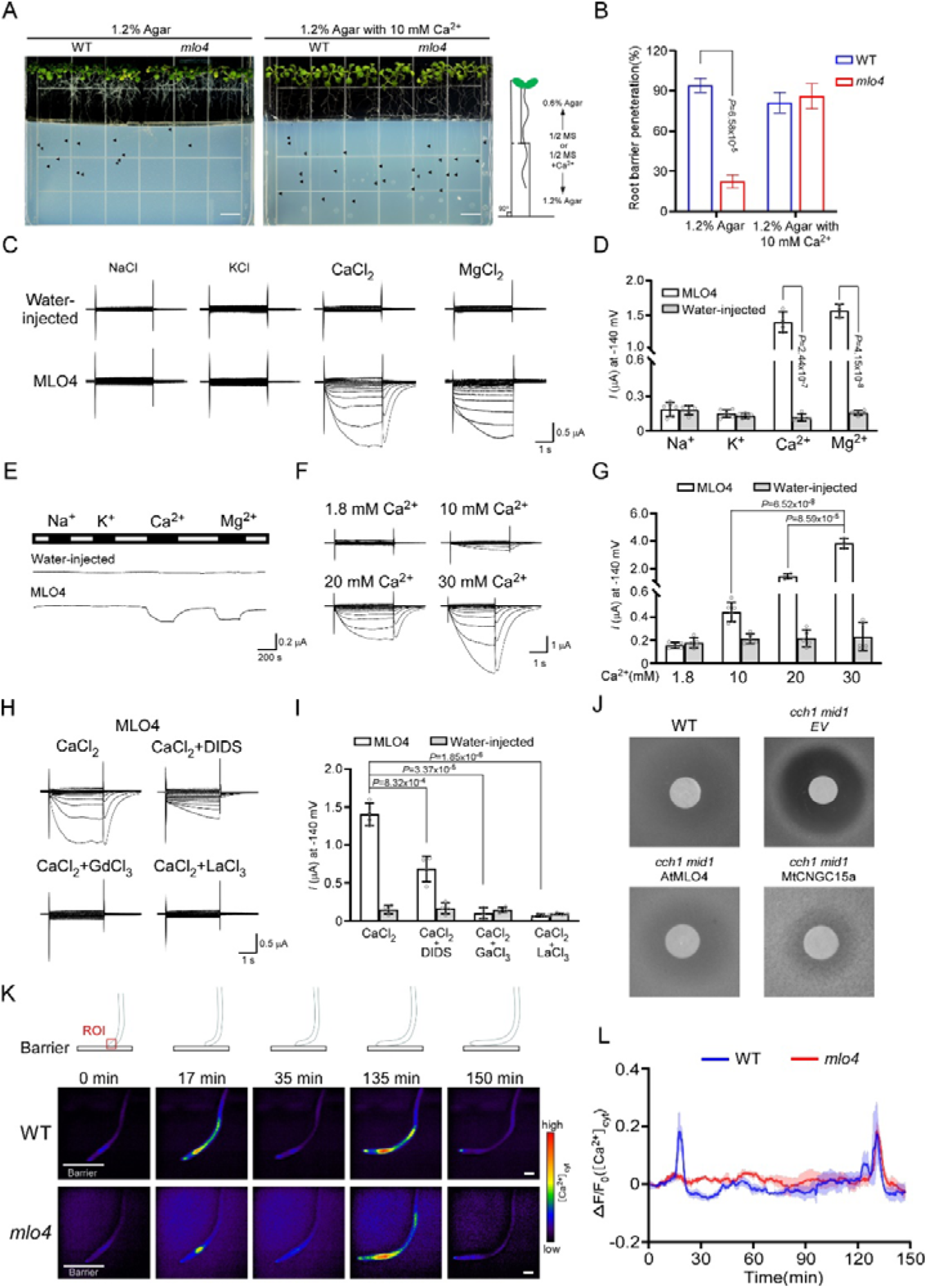
MLO4 as a calcium channel regulates root growth upon contact with hard surface of obstacles. **(A)** Penetration of roots into the hard growth media. WT and *mlo4* mutant seedlings were grown vertically on the normal growth media (1/2MS + 0.6% agar) following by reaching into the barrier (1/2MS + 1.2% agar) supplemented without (left) or with 10 mM CaCl_2_ (right). Black arrows indicate root tips within the barrier (Scale bars =7 mm). **(B)** Analysis of root penetration rates in the hard barrier for WT and *mlo4* as in (A). The data are showed as mean ± s.d., and statistical analysis is performed by two-tailed student *t*-test (WT, *n* = 4; *mlo4, n* = 4). **(C)** TEVC (Two-electrodes voltage clamp) recording from *Xenopus laevis* oocytes expressing MLO4 under different bath solutions containing 20 mM NaCl, 20 mM KCl, 20 mM CaCl_2_ and 20 mM MgCl_2_, respectively. Water-injected samples as negative controls. **(D)** Summary of steady-state currents at −140 mV from multiple recordings as in (A). The data are represented as mean ± s.d., and statistical analysis is performed by two-tailed student *t*-test (Water-injected controls, *n* = 5; NaCl, *n* = 6; KCl, *n* = 6; CaCl_2_, *n* = 4; MgCl_2_, *n* = 3). **(E)** Gap-free recordings from *Xenopus laevis* oocytes expressing MLO4 under different bath solutions containing 20 mM NaCl, 20 mM KCl, 20 mM CaCl_2_ and 20 mM MgCl_2_, respectively. Water-injected samples as negative controls. **(F)** TEVC recording from *Xenopus laevis* oocytes expressing MLO4 under different bath solutions containing 1.8 mM CaCl_2_, 10 mM CaCl_2_, 20 mM CaCl_2_ and 30 mM CaCl_2_,, respectively. **(G)** Summary of steady-state currents at −140 mV from multiple recordings as in (A). The data are represented as mean ± s.d., and statistical analysis is performed by two-tailed student *t*-test (Water-injected controls, *n* = 5; bath solution, *n* = 5; 10 mM CaCl_2_, *n* = 6; 20 mM CaCl_2_, *n* = 4; 30 mM CaCl_2_, *n* = 3). **(H)** TEVC recording from *Xenopus laevis* oocytes expressing MLO4 under different bath solution containing 20 mM CaCl_2_, 20 mM CaCl_2_ + 100μM DIDS, 20 mM CaCl_2_ + 50 μM GdCl_3_, 20 mM CaCl_2_ + 50 μM LaCl_3_. **(I)** Summary of steady-state currents at −140 mV from multiple recordings as in (C). The data are represented as mean ± s.d., and statistical analysis is performed by two-tailed student *t*-test (Water-injected controls, *n* = 4; CaCl_2_, *n* = 4; CaCl_2_ + DIDS, *n* = 4; CaCl_2_ + GdCl_3_, *n* = 3; CaCl_2_ + LaCl_3_, *n* = 4). **(J)** Measurement of MLO4 activity in the complementation assay in yeast. Like the positive control MtCNGC15, MLO4 can also complement the mutant strain *cch1/mid1* to grow under the treatment of inhibitor α-factor. WT and *cch1/mid1* transformed by empty vector are positive and negative controls, respectively. **(K)** Time-course analysis of mechanical stimuli induced increases in [Ca^2+^]_cyt_. WT and *mlo4* mutant both stably express Ca^2+^ indicator GCaMP6s. The red rectangle is marked as the region of interest (ROI) for Ca^2+^ intensity measurement, Scale bars =50 μm). **(L)** Summary of [Ca^2+^]_cyt_ signal responses after mechanical stimuli in root tips of WT and *mlo4* (mean ± s.e.m., *n* = 3). The first spike is disappeared in *mol4*.

To test the hypothesis that MLO4 may function as a calcium channel, we cloned the full-length coding sequence of *MLO4* into oocyte expression vector and examined the channel activity of MLO4 in oocytes using the two-electrode voltage-clamp (TEVC) method. When Ca^2+^ was added as the sole cation in the bath solution, TEVC recording showed that the oocytes injected with *MLO4* cRNA (complementary RNA made *in vitro* from cDNA) displayed large inward currents of 1400 ± 150 nA at −140 mV with unique tail currents in the presence of 20 mM CaCl_2_ (Figure 1C-D). To determine whether the current resulted from Ca^2+^ or chloride, we replaced CaCl_2_ with Ca(Glu)_2_ in the bath solution and again recorded the similar current, suggesting that calcium may be the charge carrier (Supplementary Figure 2). When perfused with 20 mM MgCl_2_, the oocytes expressing *MLO4* generated large currents of 1500 ± 90 nA at the same holding potential without the tail currents (Figure 1C-D). The fact that Ca^2+^, but not Mg^2+^, produced a unique tail current suggested that this tail current may be from CaCC (calcium-activated chloride channel) of oocytes [6,7]. Next, we used “gap-free” approach that recorded the same cell at the same hold potential of −70 mV while sequentially changing bath solution containing 20 mM KCl, NaCl, CaCl_2_ or MgCl_2_. Consistent with earlier results, a large inward current was recorded when Ca^2+^ and Mg^2+^ was in the bath of the oocyte expressing MLO4, whereas the oocytes injected with water did not produce any discernible current (Figure 1E). Meanwhile, those monovalent cations including Na^+^ and K^+^ did not elicit significant currents in the oocytes injected with *MLO4* cRNA, suggesting that MLO4 is selectively permeable to divalent cations such as Ca^2+^ and Mg^2+^. Furthermore, the amplitudes of recorded currents were enhanced as the concentration of Ca^2+^ increased from 10 mM to 20 mM and 30 mM, indicating that the currents depend on the Ca^2+^ concentrations at hyperpolarization pulses (Figure 1F-G).

It has been reported that CaCC is present in the *X. laevis* oocytes [6,7], which in this case would be activated by MLO4-mediated calcium influx. Therefore, the total inward current produced in the presence of 20 mM CaCl_2_ in the bath solution consisted of both CaCC-mediated chloride current and Ca^2+^-influx current. In the presence of MgCl_2_ in the bath, the current did not have CaCC contribution and showed different current pattern without any tail currents (Figure 1H). These features are reminiscent to those found with several Ca^2+^-permeable cation channels reported earlier [8]. To clarify this point, we added 4, 4′diisothiocyanostilbene-2, 2′-disulfonic acid (DIDS), a chloride channel blocker that inhibits CaCC activity. When 100 μM DIDS was included in the bath solution containing CaCl_2_, the inward current mediated by CaCC largely disappeared as reflected by removal of “tail” current. If both Ca^2+^-currents and CaCC currents depend on Ca^2+^ influx mediated by MLO4, all inward current should be blocked by the typic Ca^2+^ channel inhibitors like GdCl_3_ or LaCl_3_. Indeed, the inward currents produced by MLO4 in the presence of 20 mM CaCl_2_ disappeared when we added LaCl_3_ or GaCl_3_ (Figure 1H). Taken together, these results indicate that MLO4 constitutes a novel Ca^2+^-permeable channel.

We also used another model system, yeast *cch1/mid1* mutant, to confirm that MLO4 is a Ca^2+^ permeable channel. The *cch1/mid1* yeast mutant lacks the Cch1-Mid1 Ca^2+^ channel and fails to grow in the presence of mating pheromone in the medium [9]. This phenotype can be rescued by expression of exogenous proteins that mediate Ca^2+^ influx. Using a plant calcium channel, MtCNGC15a, as a positive control and the empty vector as negative control, we found that MLO4, like MtCNGC15a, rescued the *cch1/mid1* from growth arrest by the pheromone, confirming that MLO4 mediated Ca^2+^ influx into yeast cells (Figure 1J).

Previous studies show that cytoplasmic free Ca^2+^ in *Arabidopsis* roots changes in response to touch [1,2,10–12]. Since the MLO4 protein served as a Ca^2+^ influx channel and disruption of its expression caused defect in root mechanosensing, we hypothesized that MLO4 may be one of the channels responsible for producing touch-sensitive Ca^2+^ signals. To test this idea, we crossed *mlo4* mutant with the transgenic plant carrying GCaMP6s calcium indicator to produce *mlo4/GCaMP6s*. We then examined the Ca^2+^ spikes in 7-day seedlings in response to a hard surface during root elongation. The root tip cells in the wild-type plants showed increased [Ca^2+^]_cyt_ at 17 min after reaching the barrier. The root continued to grow along the surface of the barrier and a second Ca^2+^ spike appeared at 135 min. However, the first Ca^2+^ spike was hardly detectable in *mlo4* root tip cells although the second Ca^2+^ spike was similar to that in the wild type during the extended growth along the barrier (Figure 1K-L). We then produced plants expressing a FRET calcium indicator *YC3.6*. Comparing wild type and mutant carrying *YC3.6*, we found that the touch-induced Ca^2+^ signal in wild type plants was essentially diminished in the *mlo4/YC3.6*. Taken together, these results supported the conclusion that MLO4 is required for producing mechanically triggered [Ca^2+^]_cyt_ elevation in root tip cells (Supplementary Figure 3).

In summary, we reported here that a MLO family member, MLO4, functions as a typical Ca^2+^ channel that links touch stimulation to Ca^2+^ elevation in root tip cells. Future studies will be directed to expand the possibility that all MLO family proteins may function as calcium channels that play essential roles in a variety of physiological processes, including mildew resistance in which the MLOs were originally identified. Further work will also identify the regulatory mechanisms underlying activation of the MLO-type channels and how these channel work together with other channels to generate a large array of Ca^2+^ signatures in plant cells.

---

**Figure S1.**
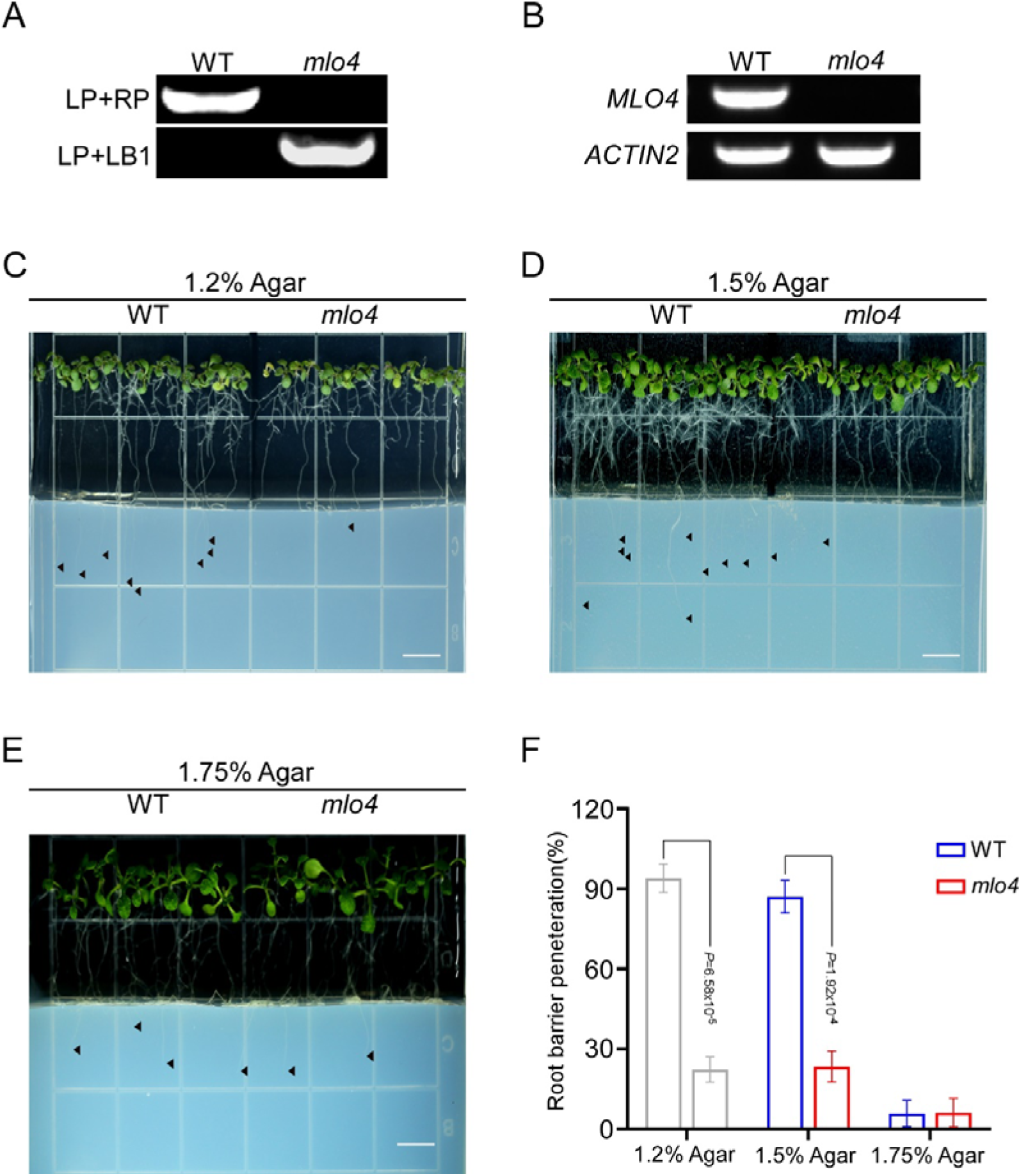
The root penetration into the hard growth media is repressed significantly in *mlo4* mutants. **(A-B)** Identification of T-DNA insertion mutant of *mlo4* in DNA level (A) and RNA level (B). **(C-E**) Penetration of roots into the hard growth media. WT and *mlo4* mutant seedlings were grown vertically on the normal growth media (1/2MS + 0.6% agar) following by reaching into the barrier with different concentrations of agar (1/2MS + 1.2% agar1.5%, or 1.75% agar, respectively). Black arrows indicate root tips within the barrier (Scale bars =7 mm). **(F)** Analysis of root penetration rates in the hard barrier for WT and *mlo4* as in (C-E). The data are showed as mean ± s.d., and statistical analysis is performed by two-tailed student *t*-test (WT, *n* = 4; *mlo4, n* = 4).

**Figure S2.**
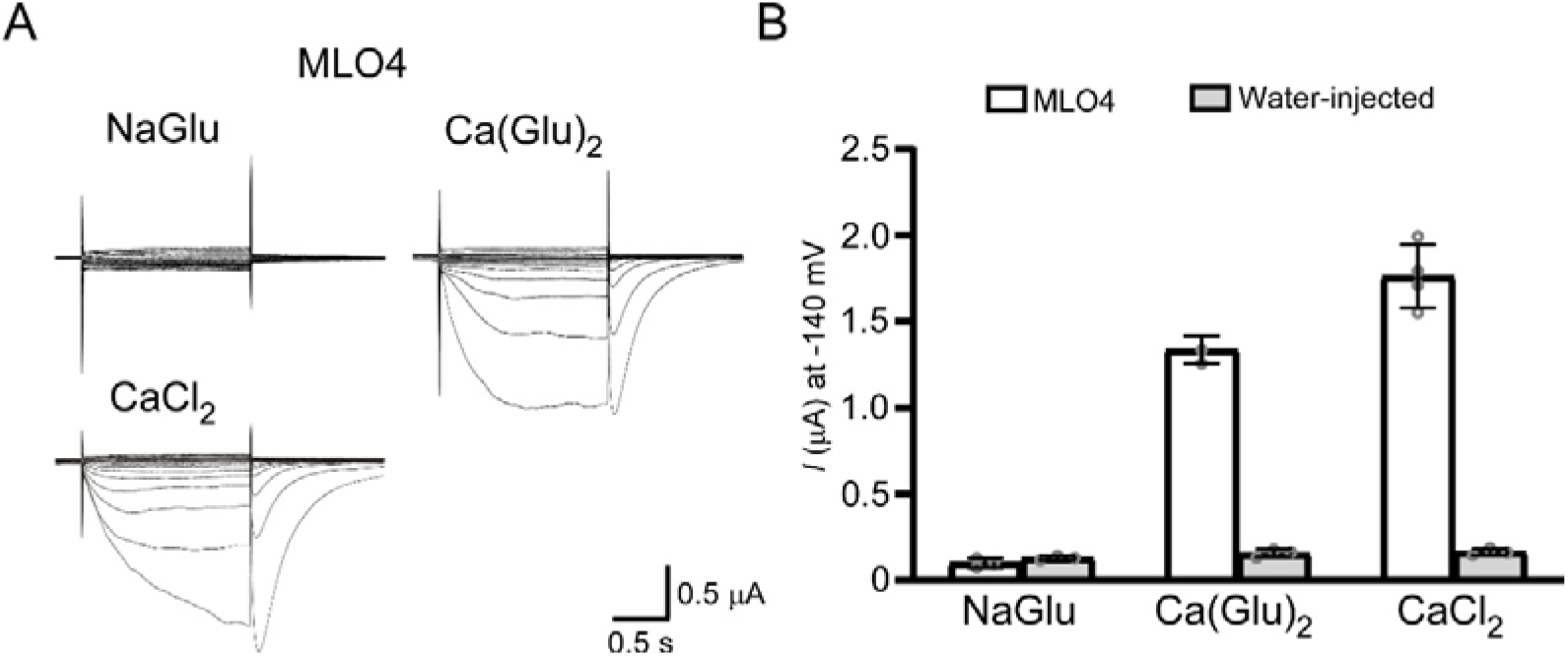
The currents mediated by MLO4 depend on Ca^2+^. **(A)** TEVC recording from *Xenopus laevis* oocytes expressing MLO4 under different bath solutions containing 20 mM NaGlu, 20 mM Ca(Glu)_2_, 20 mM CaCl_2_, respectively. **(B)** The data are represented as mean ± s.d., and statistical analysis is performed by two-tailed student *t*-test (Water-injected controls, *n* = 3; NaGlu, *n* = 3; Ca(Glu)_2_, *n* = 3; CaCl_2_, *n* = 4).

**Figure S3.**
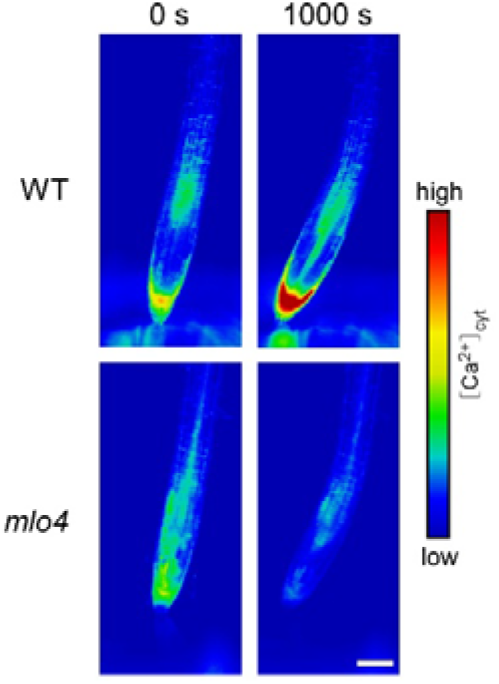
Mechanical stimuli induced [Ca^+^]_cyt_ elevation is disappeared in *mlo4* root tips. Time-course analysis of mechanical stimuli induced increases in [Ca^2+^]_cyt_. WT and *mlo4* mutant stably express fluorescence resonance energy transfer (FRET)-based Ca^2+^ sensor YC3.6. Time-lapse emission ratio images are shown using a pseudo-color (Scale bars = 50 μm).

## AUTHOR CONTRIBUTIONS

C.H., Z.Z., S.L., L.L. conceived and designed the project. Z.Z., Q.N. conducted the electrophysiological experiments; Y.S. performed molecular cloning and yeast complementation; F.C., P.L., G.L. conducted total internal reflection fluorescence experiments; P.L., G.L. performed transgenic plant generation and the growth phenotype analysis. All experiments were independently reproduced in the laboratory. L.L., Z.Z., C.H. analyzed the data and wrote the manuscript. S.L. revised the manuscript. All authors discussed the results and commented on the manuscript.

## ACKNOWLEDGEMENTS

This work was supported by grants from the Key Program of the National Natural Science Foundation of China (31930010 to L.L.), the General Program of National Natural Science Foundation of China (No. 31872170 to L.L. and No. 31900234 to C.H.).

